# Dopamine receptor 1 on CaMKII-positive neurons within claustrum mediates adolescent cocaine exposure-induced anxiety-like behaviors and electro-acupuncture therapy

**DOI:** 10.1101/2022.10.18.512685

**Authors:** Liying Chen, Zhaoyu Liu, Ziheng Zhao, Demin Du, Weichao Pan, Xiaoyan Wei, Jiaxun Nie, Feifei Ge, Jianhua Ding, Yu Fan, Hee Young Kim, Xiaowei Guan

## Abstract

Adolescent cocaine exposure (ACE) increases risk of developing psychiatric problems such as anxiety, which may drive relapse in later life, however, its underlying molecular mechanism remains poorly understood.

**Methods:** ACE male mice model were established by exposing to cocaine during adolescent period. Elevated plus maze (EPM) were used to assess anxiety-like behaviors in mice. Within claustrum, local injection of SCH-23390, a specific antagonist for dopamine receptor 1 (D1R), or D1R knocking-down virus were used to regulate D1R function or expression on CaMKII-positive neurons (D1R^CaMKII^) *in vivo*. Electro-acupuncture (EA) treatment was performed at acupoints of Baihui and Yintang during withdrawal period.

**Results:** We found that ACE mice exhibited anxiety-like behaviors, along with more activated CaMKII-positive neurons and increased D1R^CaMKII^ levels in claustrum during adulthood. Inhibiting D1R function or knocking-down D1R^CaMKII^ levels in claustrum efficiently reduced claustrum activation, and ultimately suppressed anxiety-like behaviors in ACE mice during adulthood. EA treatment alleviated ACE-evoked claustrum activation and anxiety-like behaviors by suppressing claustrum D1R^CaMKII^.

**Conclusion:** Our findings identified a novel role of claustrum in ACE-induced anxiety-like behaviors, and put new insight into the D1R^CaMKII^ in the claustrum. The claustrum D1R^CaMKII^ might be a promising pharmacological target, such as EA treatment, to treat drugs-induced anxiety-like behaviors.

**Graphic abstract:** 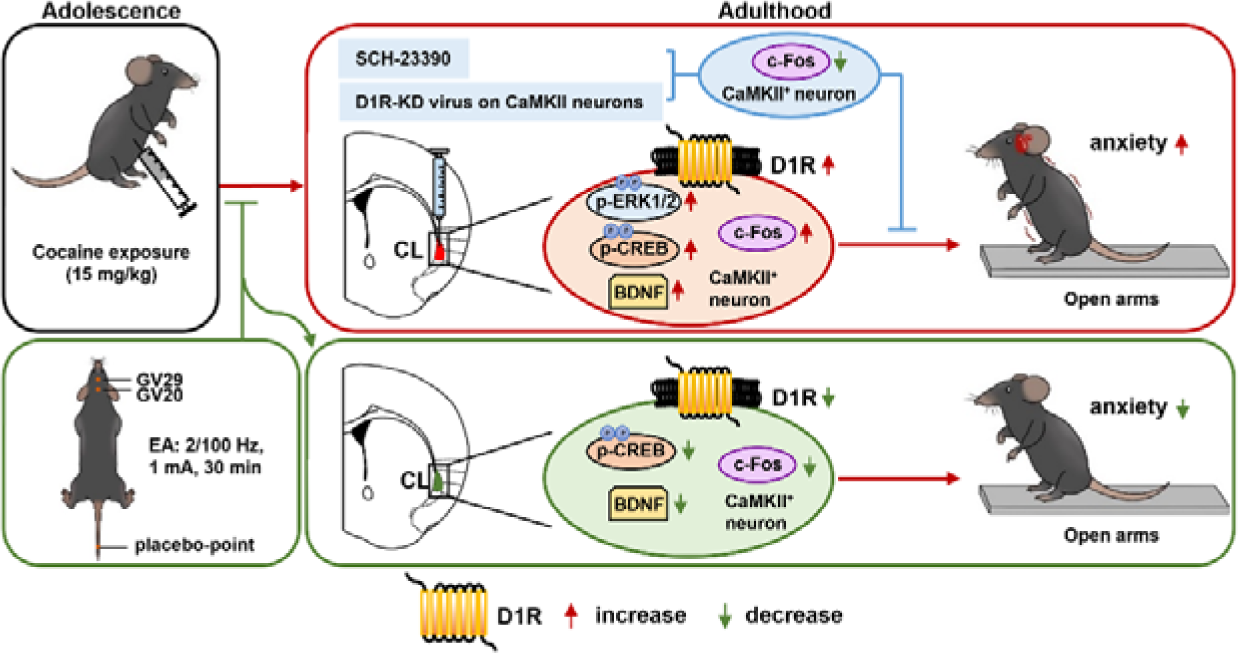

## Introduction

Adolescents present higher vulnerability to substance abuse [1,2]. Adolescent experience of drugs increase the risk of developing psychiatric disorders that even persists into adulthood [3,4]. Anxiety is a prominent psychiatric symptom of adolescent drug abuse, which is generally believed as critical factor for driving relapse to drugs in later life [5,6]. Previously, we showed that adolescent cocaine-exposed (ACE) mice exhibit more anxiety-like behaviors in their adulthood [7,8], however, the involved brain region remains much to be clarified.

The claustrum is a thin stripped nucleus located between insular cortex and putamen. Above 90% neurons in claustrum belong to CaMKII-positive glutamatergic neurons [9]. Anatomically, the claustrum has extensive reciprocal connectivity with most neocortical regions [10–13] and communicates densely with limbic structure and thalamus [14,15], implying that claustrum may be a hub for integrating and relaying “executive process” of cortex. At present, the functions of claustrum remain poorly understood. Emerging studies demonstrate that claustrum may be key nucleus involved in consciousness [16], attention [17,18] and cognition [19–21]. The claustrum has been implicated in numerous brain disorders, such as seizures [22], autism [23] and Parkinson Disease [24], but it has not been evidently as the core structure of any specific disease. In marijuana users [25,26] or internet game addicts [27], the claustrum shows greater activity. Recently, Terem et al. [28] found that the dopamine receptor 1 (D1R)-positive neurons in claustrum is required for acquisition of cocaine-preferred behaviors, and stimulating these neurons could drive context-reinstatement of cocaine preference. In addition, the other recent study shows that the claustrum mediates stress-induced anxiety in mice [29]. These studies demonstrate that claustrum is involved in addiction and anxiety, but its role in drugs-related anxiety is unknown.

Currently, there is few pharmacotherapy options for adolescent substance use-induced anxiety, partially due to serious side-effects caused by anxiolytic medicine. As a non-invasive therapeutic approach, acupuncture has been considered as an efficient intervention on substance use, especially for treatment of the protract abstinence syndrome (such as anxiety) [30,31], but its pharmacological targets remain largely unclear. The electro-acupuncture (EA), a kind of peripheral acupuncture with electric stimulation, has been shown to improve anxiety without adverse effects [32,33]. Previously, we found that EA at acupoints of Baihui (GV 20) and Yintang (GV 29) effectively reduced the anxiety-like behaviors in ACE mice during adulthood [34]. Early in 2006, Shapo et al. [35] reported that acupuncture at Weizhong points (BL40) could significantly evoke claustrum in humans. Whether claustrum is one of key targets for EA to treat ACE-related anxiety, and its potential pharmacological mechanism, has not been explored yet.

In the present study, male mice were exposed to cocaine during adolescent period from postnatal day 28 (P28) to P42 [36]. EA treatment at acupoints of Baihui (GV20) and Yintang (GV29) were performed in male mice during withdrawal period. When growing up to adult ages, the mice were subjected to anxiety behavioral tests to explore the role of claustral D1R, especially expressed on CaMKII-positive neurons (D1R^CaMKII^), in ACE-induced anxiety-like behaviors and the EA therapy.

## Materials and methods

### Animals

Male C57BL/6 wild type (WT) mice of postnatal day 21 (P21), weighing 12-16 g were used. All animals were housed at constant humidity (40∼60%) and temperature (24 ± 2°C) with a 12-hour light/dark cycle (lights on at 8 a.m.) and allowed free access to food and water. Handling of the animals was carried out for three days before onset of experiments. All procedures were carried out in accordance with the National Institutes of Health Guide for the Care and Use of Laboratory Animals and approved by the Institutional Animal Care and Use Committee (IACUC) at Nanjing University of Chinese Medicine, China.

### Drug treatment

On P28, male C57BL/6 WT mice were exposed to cocaine hydrochloride (15 mg/kg, dissolved in saline, i.p., Qinghai Pharmaceutical, China) or saline (0.2 mL, i.p.) once daily at 10 a.m. for 14 consecutive days. From P42-P70, these mice were kept at their homecage (four mice per cage). On P71, elevated plus maze (EPM) tests were performed in mice. To exclude the influence of EPM procedure on claustrum activity and related molecules expression, the brain tissue was collected on P77 for qPCR, western blot and immunofluorescence.

### Electro-acupuncture treatment

During P42 to P70, two acupuncture needles (0.16 × 7 mm, Hwato, China) were inserted gently in a depth of 2–3 mm at the acupoints of Baihui (GV20, the midpoint between the auricular apices) and Yintang (GV29, the midpoint between the medial ends of the two eyebrows) in each mouse, the acupoints were electric mixed stimuli of 2/100 Hz at an intensity of 1 mA (HANS Electronic Apparatus, China) for 30 min once daily at acupoints of GV20 and GV29 in EA mice, or at “placebo-point” (at 1 cm from base of the tail) in Sham EA (SEA) mice.

### Behavioral test

On P71, EPM was performed to assess anxiety-like behaviors in mice (n = 119 mice). All behavioral tests were conducted during light cycle. The EPM apparatus (TopScan, USA) consists of 4 elevated arms (52 cm above the floor) arranged in a cross pattern with two 1-cm walled arms (open arms) and two 40-cm walled arms (closed arms). Each mouse was placed in the center portion of the EPM with its head facing an open arm and allowed freely to explore the maze for 5 min. The spending time in the open arms, the entries into the open arms, and the total distance traveled by mice in the EPM apparatus were recorded.

### Western blot

On P77, mice brains (n = 24 mice) were rapidly removed following the anesthesia and placed in O.C.T. compound (SAKURA, Japan). The sample were frozen quickly in liquid nitrogen and fixed on the cold sample holder. The brain tissue was coronally cut by Leica CM1950 cryostat (Leica, Germany). It was determined that Bregma +0.62 mm lever when the connected corpus callosum appeared on both sides of the sample section and then continue cutting 500 μm. A handled biopsy punch (Miltex Instruments, Japan) vertically inserted into the outside of the end of the ventrolateral corpus callosum, and claustrum tissue were collected and frozen on dry ice. The total protein was extracted from claustrum in mice using RIPA lysis buffer (Beijing ComWin Biotech Co., China) according to the manufacturer’s instructions. Protein samples (15 μg) was separated by 10% SDS–PAGE, 12% SDS–PAGE and electrophoretically transferred onto PVDF membranes. The transferred membranes were blocked with 5% non-fat dry milk or 5% BCA and 0.1% Tween 20 in 10 mM Tris–HCl (TBST buffer) for 1.5 h at room temperature, then subsequently incubated with the following primary antibodies: D1R (1:1000, Rabbit, RRID:AB_10598308, Proteintech, USA), D2R (1:1000, Rabbit, ER1907-70, Hua An Biotechnology Company, China), D3R (1:1000, Mouse, ab155098, Abcam, UK), ERK_1/2_ (1:1000, Rabbit, BM4326, Boster, China), p-ERK_1/2_ (1:1000, Rabbit, BM4326, Boster, China), CREB (1:1000, Rabbit, RRID:AB_10699020, Cell Signaling Technology, USA), p-CREB (1:1000, Rabbit, RRID:AB_2561044, Cell Signaling Technology, USA), c-Fos (1:2000, Rabbit, RRID:AB_2247211, Cell Signaling Technology, USA), BDNF (1:1000, Mouse, RRID:AB_2881675, Proteintech, USA). The next day, the membranes were washed three times in Tris-buffered saline with Tween 20 and incubated with horseradish peroxidase (HRP)-conjugated secondary antibody goat anti-rabbit or goat anti mouse (1:5000, Beijing ComWin Biotech Co., China) at room temperature for 1 h. The blots were visualized by the ECL kit (Beyotime, China) and the signal was visualized by imaging system (Tanon-5200, China). The blots were washed with stripping buffer (Beyotime Institute of Biotechnology, China) to reprobe with other antibodies. In this study, GAPDH or β-actin was used as the loading control. Values for target protein levels were calculated using Image J software (NIH, USA). The relative level of each protein expression (blot volume) was normalized to GAPDH or β-actin.

### RNA extraction and quantitative real-time PCR

On P77, mice brains (n = 6 mice) were collected and total RNA was extracted from claustrum using FastPure Cell/Tissue Total RNA Isolation Kit (Vazyme, China), according to the manufacturer’s instructions. All samples were tested for concentration and purified using a 4200 TapeStation (Agilent Technologies, Santa Clara, USA). After RNA extraction, the amount of RNA was normalized across samples, and cDNA was created using iScript (1708891, Bio-Rad, USA). The *actin* was used as the internal control. The primers of *d1r, d2r*, *d3r* and *actin* were as follows: *d1r* (forward, ATGGCTCCTAACACTTCTACCA; reverse, GGGTATTCCCTAAGAGAGTGGAC); d2r (forward, ACCTGTCCTGGTACGATGATG; reverse, GCATGGCATAGTAGTTGTAGTGG); *d3r* (forward AGACAACATGGAGAGCTGAAACGCT; reverse, TTCAGGGCTGTGGATAACCTGCC); *actin* (forward, GGCTGTATTCCCCTCCATCG; reverse, CCAGTTGGTAACAATGCCATGT). *d1r*, *d2r* and *d3r* mRNA levels were normalized to *actin* mRNA levels. The relative mRNA level was calculated by the comparative CT method (2^-ΔΔCt^).

### Immunofluorescence

On P77, mice brains (n = 70 mice) were collected and perfused with 4% paraformaldehyde (PFA). The coronal brain sections (30 μm) were cut on a cryostat (Leica, Germany). The sections were incubated in 0.3% (v/v) Triton X-100 for 0.5 h, blocked with 5% donkey serum for 1.5 h at room temperature, and incubated overnight at 4°C with the following primary antibodies: rabbit anti-c-Fos (1:2000, RRID: AB_2247211, Cell Signalling Technology, USA), mouse anti-CaMKIIα (1:100, RRID: AB_626789, Santa Cruz, USA) and rabbit anti-D1R (1:100, RRID: AB_10598308, Proteintech, USA), followed by the corresponding fluorophore-conjugated secondary antibodies for 1 h at room temperature. The following secondary antibodies were used here: Alexa Fluor 555-labeled donkey anti-rabbit secondary antibody (1:500, RRID: AB_162543, Invitrogen, USA), Alexa Fluor 488-labeled donkey anti-mouse secondary antibody (1:500, RRID: AB_141607, Invitrogen, USA). Fluorescence signals were visualized using a Leica DM6B upright digital research microscope (Leica, Germany) or Leica TCS SP8 (Leica, Germany) with the parameters including Format 1024×1024, Scan Speed 200 Hz, Zoom 1. The immunofluorescent signal values of D1R receptors on CaMKII-positive neurons in the claustrum region of each slice (total value) were calculated by LAS X software (version: 3.5.6.21481). The mean gray value of D1R on each CaMKII-positive neuron were obtained by total value of D1R divide the number of the CaMKII-positive neuron in one slice. We calculated at least 3 slices in one mouse, and 6 mice in one group.

### Local injection of SCH-23390 in claustrum

The D1 receptor antagonist R(+)-7-chloro-8-hydroxy-3-methyl-1-phenyl-2,3,4,5-tetrahydro-1H-3-benzazepine hydrochloride (SCH-23390, Selleck chemicals, China) was dissolved in 0.9% saline and stored -80°C. On P64, the cannula (O.D. 0.41 mm, C.C 3.0 mm, Cat 62004, RWD, China) were implanted into claustrum (AP + 0.62 mm, ML ± 3.05 mm, DV – 2.50 mm) in mice brain (n = 30 mice). After surgery, mice were maintained at homecage about 1 week. On P71, a volume of 70 nL of saline (vehicle, Veh) or SCH-23390 (SCH) were bilaterally injected into claustrum at a rate of 10 nL/min at 20 min prior to EPM behavioral tests. On P77, a volume of 70 nL of saline (vehicle) or SCH-23390 were bilaterally injected into claustrum at a rate of 10 nL/min at 20 min prior to brain tissue collection.

### Electrophysiology

The naïve mice (n = 4 mice) were deeply anesthetized with isoflurane (RWD, China) and perfused with ice-cold cutting solution. Slice preparation was performed as previously described [37]. Slices containing the claustrum were cut at a 200 μm thickness using a vibratome in 4°C cutting solution. The slices were transferred to 37°C cutting solution for 9 min, then transferred to a holding solution to allow for recovery at room temperature for 1 h before recordings. During electrophysiological recordings, the brain slice was continuously perfused with oxygenated artificial CSF (aCSF) maintained at 30°C by a solution heater (TC-324C, Warner Instruments, USA).

Recordings were performed under current-clamp mode. Whole-cell current-clamp microelectrodes (3–5 MΩ) were filled with potassium internal solution. For measurements of neuronal excitability, current-step protocols were run (step, 20 pA; action potential range, 0–200 pA) at -70 mV. Action potential (AP) signals were filtered at 4 kHz, amplified at 5× using a MultiClamp 700B amplifier (Molecular Devices, USA) and digitized at 10 kHz with a Digidata 1440A analog-to-digital converter (Molecular Devices, USA).

The AP baseline (pre-SCH-23390 application) was recorded for 5 min. The SCH-23390 (10 μM) was then added to aCSF and the brain slices were bathed for 10 min before the same protocol was repeated. The number of AP in each cell was analyzed with Clampfit 10.6 software (Molecular Devices, USA) in response to each current step in the presence of SCH-23390 compared to the baseline.

### Local knockdown of D1R on CaMKII-positive neurons of claustrum

On P50, mice (n = 34 mice) were anesthetized with 2% isoflurane in oxygen, and were fixed in a stereotactic frame (RWD, China). A heating pad was used to maintain the core body temperature of the mice at 36°C. The coordinates of claustrum were defined as AP + 0.62 mm, ML ± 3.05 mm, DV –3.50 mm. A volume of 70 nL virus of *CaMKIIap-EGFP-MIR155(shDrd1)-SV40 PolyA* (KD group, Titer: 8.89E+13v.g/ml, GeneChem, China) or *CaMKIIap-EGFP-MIR155(MCS)-SV40 PolyA* (Ctrl group, Titer: 1E+13v.g/ml, GeneChem, China) were injected bilaterally into claustrum at a rate of 10 nL/min. After surgery, mice were kept at homecage about 3 weeks.

### Statistical analysis

Statistical analysis was carried out using GraphPad Prism software (Version 8.0.2). Data are presented as the Mean ± SEM. The behavioral data and electrophysiological data were analyzed by Two-way analysis of variance (ANOVA) with Sidak’s multiple comparisons, other data was analyzed by unpaired *t-test*. Statistical significance was set as *p* < 0.05.

## Results

### ACE evokes CaMKII-positive neurons and D1R^CaMKII^ levels in claustrum, and enhances anxiety-like behaviors in mice during adulthood

To examine anxiety-like behaviors in mice, mice were subjected to behavioral tests of EPM during their adulthood (Figure 1A). As shown in Figure 1B, ACE mice (n = 16 mice) exhibited less time spent (*t* = 3.813, *p* = 0.0006) and less entries (*t* = 2.823, *p* = 0.0084) into the open arms, while similar total distances traveled (*t* = 0.6731, *p* = 0.5061) in EPM apparatus than those of adolescent saline exposure (ASE) mice (n = 16 mice) during adulthood. These results indicate that ACE cause anxiety-like behaviors in mice during adulthood.

**Figure 1.**
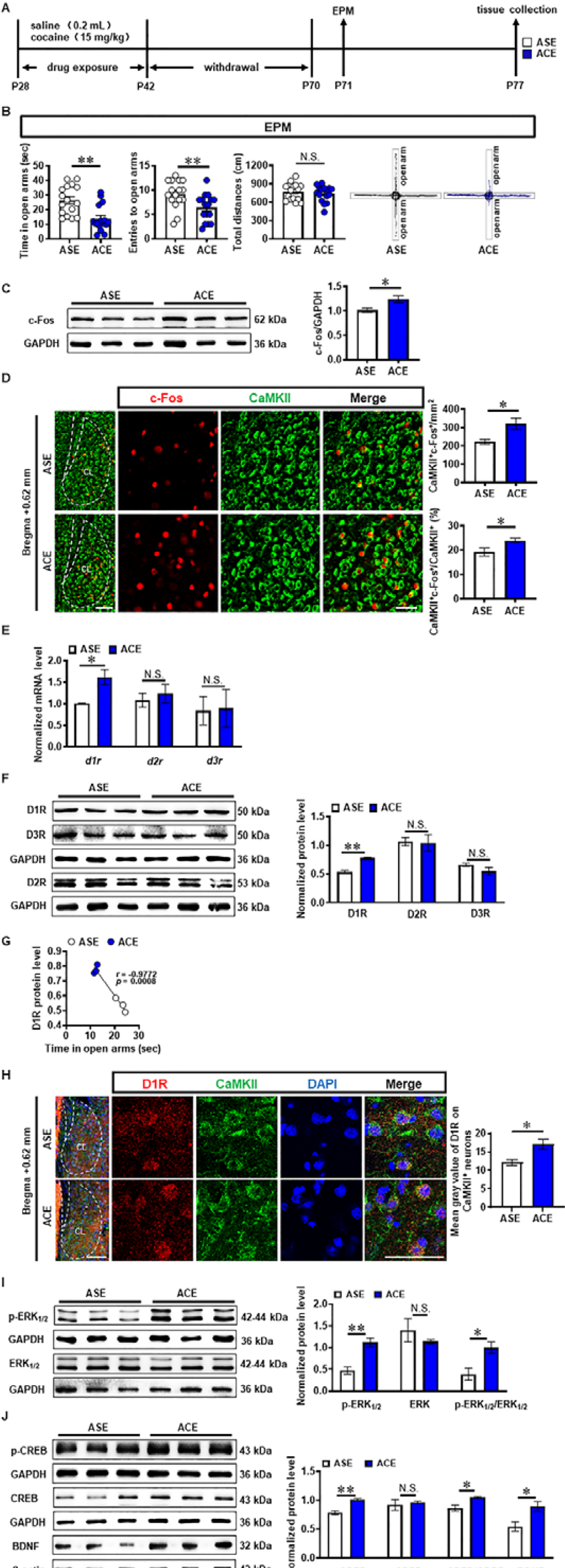
ACE enhances anxiety-like behaviors in mice during adulthood, along with the increased CaMKII-positive neuron activity and D1R^CaMKII^ level in claustrum. **A**, Experimental timeline. **B**, EPM test. n = 16 mice per group. **C**, Total protein levels of c-Fos. n = 3 mice per group. **D**, The number and percentage of c-Fos-positive & CaMKII-positive neurons. n = 6 mice per group. Scale bar, 100 μm/50 μm. **E**, The mRNA levels of *d1r, d2r and d3r*. n = 3 mice per group. **F**, The protein levels of D1R, D2R and D3R. n = 3 mice per group. **G**, The correlation analysis of claustrum D1R levels with EPM data. n = 6 mice. **H**, The expression of D1R^CaMKII^. n = 6 mice per group. Scale bar, 100 μm/50 μm. **I**, Levels of p-ERK_1/2_ and ERK_1/2._ n = 3 mice per group. **J**, Levels of p-CREB, CREB and BDNF. n = 3 mice per group. ASE, adolescent saline exposure; ACE, adolescent cocaine exposure; CL, claustrum. Data are presented as the Mean ± SEM. N.S., *p* > 0.05, *, *p* < 0.05, **, *p* < 0.01 vs ASE.

The c-Fos and CaMKII were used as markers to detect the neuronal activity and glutamatergic neurons, respectively. The total protein level of c-Fos (*t* = 2.811, *p* = 0.0483) was higher in ACE mice (n = 3 mice) than that in ASE mice (n = 3 mice) during adulthood (Figure 1C). Both the number (*t* = 2.966, *p* = 0.0141) and percentage (*t* = 2.454, *p* = 0.034) of c-Fos-positive & CaMKII-positive neurons were higher in ACE mice (n = 6 mice) than those in ASE mice (n = 6 mice) during adulthood (Figure 1D). These results indicate that ACE increase claustrum activity, especially for the CaMKII-positive neurons, in male mice during adulthood.

The total levels of dopamine receptor 1 mRNA (*d1r*, *t* = 3.502, *p* = 0.0249, Figure 1E) and protein (D1R, *t* = 7.389, *p* = 0.0018, Figure 1F) of claustrum was higher in ACE mice (mRNA, n = 3 mice; protein, n = 3 mice) than those in ASE mice (mRNA, n = 3 mice; protein, n = 3 mice) during adulthood. There were no significant differences in claustrum dopamine receptor 2 mRNA (*d2r*, *t* = 0.5756, *p* = 0.5957, Figure 1E) and protein (D2R, *t* = 0.1621, *p* = 0.8791, Figure 1F), and claustrum dopamine receptor 3 mRNA (*d3r*, *t* = 0.1051, *p* = 0.9214, Figure 1E) and protein (D3R, *t* = 1.519, *p* = 0.2035, Figure 1F) between ACE mice (mRNA, n = 3 mice; protein, n = 3 mice) and ASE mice (mRNA, n = 3 mice; protein, n = 3 mice) during adulthood. As shown in Figure 1G, the claustrum D1R level were negatively correlated with the spending time in open arms of EPM apparatus in mice during adulthood (n = 6 mice, *r* = -0.9772, *p* = 0.0008). On CaMKII-positive neurons, the mean gray value of D1R was much higher (*t* = 3.062, *p* = 0.0120) in ACE mice (n = 6 mice) than that in ASE mice (n = 6 mice) during adulthood (Figure 1H). These results indicate that ACE obviously increase *d1r* and D1R level, particularly those expressed on CaMKII-positive neurons in claustrum during adulthood.

In parallel, the protein levels of p-ERK_1/2_ (*t* = 4.905, *p* = 0.0080), ratio of p-ERK_1/2_ to ERK_1/2_ (*t* = 3.239, *p* = 0.0317), p-CREB (*t* = 5.871, *p* = 0.0042), ratio of p-CREB to CREB (*t* = 3.502, *p* = 0.0248) and BDNF (*t* = 2.940, *p* = 0.0424) were higher in ACE mice (n = 3 mice) than those in ASE mice (n = 3 mice), but those of ERK_1/2_ (*t* = 0.9475, *p* = 0.3970) and CREB (*t* = 0.4336, *p* = 0.6869) had no significant difference between ACE (n = 3 mice) and ASE mice (n = 3 mice) during adulthood (Figure 1I, 1J). These results indicate that ACE increase the potential subsequent signals of D1R^CaMKII^ in claustrum, which may be associated with ACE-induced claustrum activation and anxiety-like behaviors during adulthood.

### Locally inhibiting claustrum D1R suppresses ACE-induced anxiety-like behaviors in mice during adulthood and claustrum activation

To evaluate the role of claustrum D1R in ACE-induced anxiety-like behaviors in mice during adulthood, SCH-23390, a selective D1R antagonist, was bilaterally injected into claustrum 20 min prior to EPM tests on P71 (Figure 2A). Moreover, to exclude the potential influence of EPM procedure on claustrum activity and related molecules expression, the brain tissue was collected on P77, at 20 min after SCH-23390 injection.

**Figure 2.**
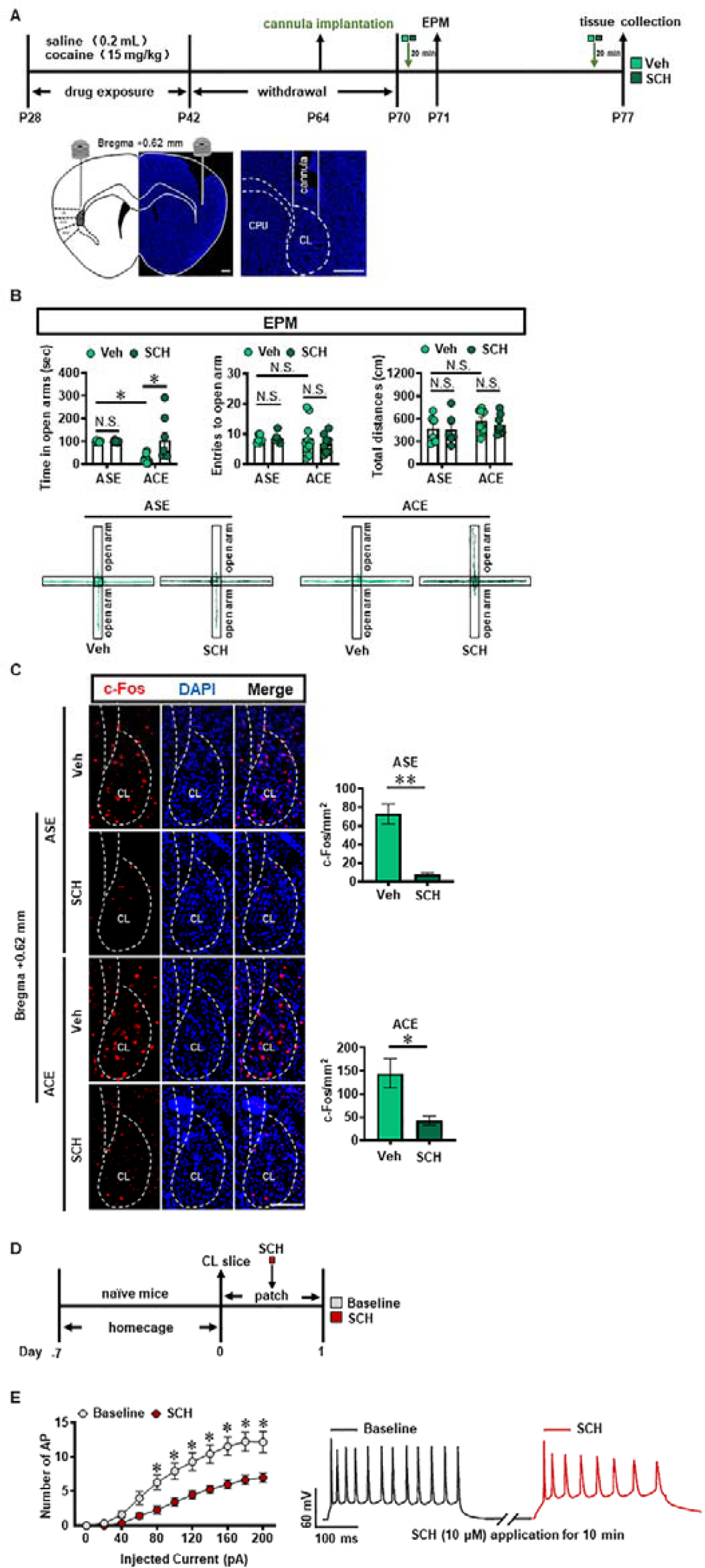
Inhibiting claustrum D1R suppresses ACE-induced anxiety-like behaviors in mice during adulthood and claustrum activation. **A**, Experimental timeline for SCH-23390 treatment and diagram of cannula implantation in claustrum. Scale bar, 400 μm. **B**, EPM test. Vehicle-treated ASE mice, n = 6 mice; SCH-23390-treated ASE mice, n = 6 mice; Vehicle-treated ACE mice, n = 10 mice; SCH-23390-treated ACE mice, n = 8 mice. **C**, The number of c-Fos-positive neurons in claustrum. n = 6 mice per group. Scale bar, 100 μm. **D**, Experimental timeline for whole-cell patch in claustrum slices. **E**, Action potentials (AP) of claustrum neurons under whole-cell current-clamp configuration (n = 16 cells from 4 mice) and sample traces for the number of AP following 200 pA current injection in the absence and presence of SCH-23390 on the same cell. Veh, vehicle-treated mice; SCH, SCH-23390-treated mice; CL, claustrum. Data are presented as the Mean ± SEM. N.S., *p* > 0.05, *, *p* < 0.05, **, *p* < 0.01 vs Veh or Baseline.

As shown in Figure 2B, compared with vehicle-treated ASE mice (n = 6 mice), vehicle-treated ACE mice (n = 10 mice) spent less time in open arms (*t* = 2.976, *p* = 0.0369), but had similar entries into the open arms (*t* = 0.1320, *p* > 0.9999) and the total distances traveled (*t* = 1.288, *p* = 0.7554) in EPM apparatus during adulthood. When compared to vehicle-treated ASE mice (n = 6 mice), SCH-23390-treated ASE mice (n = 6 mice) did not affect the time spent in open arms (*t* = 0.0066, *p* > 0.9999), the entries into the open arms (*t* = 0.1476, *p* > 0.9999) and the total distances traveled (*t* = 0.1345, *p* > 0.9999) in EPM apparatus during adulthood. However, compared with vehicle-treated ACE mice (n = 10 mice), SCH-23390-treated ACE mice (n = 8 mice) significantly increased the time spent in open arms (*t* = 3.423, *p* = 0.0123), but did not affect the entries into the open arms (*t* = 0.9296, *p* = 0.9320) and the total distances traveled (*t* = 0.7243, *p* = 0.9791) in EPM apparatus during adulthood. These results imply that inhibiting claustrum D1R could alleviate ACE-induced anxiety-like behaviors during adulthood.

SCH-23390 treatment significantly attenuated the number of c-Fos-positive neurons in ASE mice (n = 6 mice*, t* = 6.028, *p* = 0.0001) and ACE mice (n = 6 mice, *t* = 3.097, *p* = 0.0113), when compared with vehicle-treated ASE mice (n = 6 mice) and vehicle-treated ACE mice (n = 6 mice), respectively (Figure 2C).

To further explore the effects of D1R on the neuronal excitability in claustrum, the whole-cell recordings were performed in brain slices containing claustrum of naïve mice (Figure 2D). As shown in Figure 2E, the number of action potential (AP) was significantly reduced by SCH-23390 in claustrum slice at 80–200 pA injecting currents (F_(10,330)_ = 3.068, *p* = 0.001; 20–60 pA, *p* > 0.05; 80 pA, *p* = 0.0115; 100 pA, *p* = 0.0026; 120 pA, *p* = 0.0009; 140 pA, *p* = 0.0003; 160 pA, *p* < 0.0001; 180 pA, *p* < 0.0001; 200 pA, *p* = 0.0003) as compared to the pre-SCH-23390 baseline, indicating that SCH-23390 treatment could significantly suppress neuronal activity in claustrum slices of naïve mice.

### Knocking-down claustrum D1R^CaMKII^ level suppresses ACE-induced anxiety-like behaviors in mice during adulthood and claustrum activation

To examine the role of claustrum D1R^CaMKII^ level in the ACE-induced anxiety-like behaviors and claustrum activation, the Adeno-associated virus of either *CaMKIIap-EGFP-MIR155(shDrd1)-SV40 PolyA* (KD) or *CaMKIIap-EGFP-MIR155(MCS)-SV40 PolyA* (Ctrl) was bilaterally infused into claustrum to knocking-down the D1R^CaMKII^ in claustrum (Figure 3A). Figure 3B showed the local injection site of virus into the claustrum. As shown in Figure 3C, compared with the corresponding Ctrl-treated groups (ASE, n = 3 mice; ACE, n = 3 mice), D1R levels in claustrum were significantly reduced by KD treatment (ASE, n = 3 mice, *t* = 3.184, *p* = 0.0334; ACE, n = 3 mice, *t* = 3.465, *p* = 0.0257), indicating an efficient knockdown effect of KD virus on D1R^CaMKII^ in claustrum.

**Figure 3.**
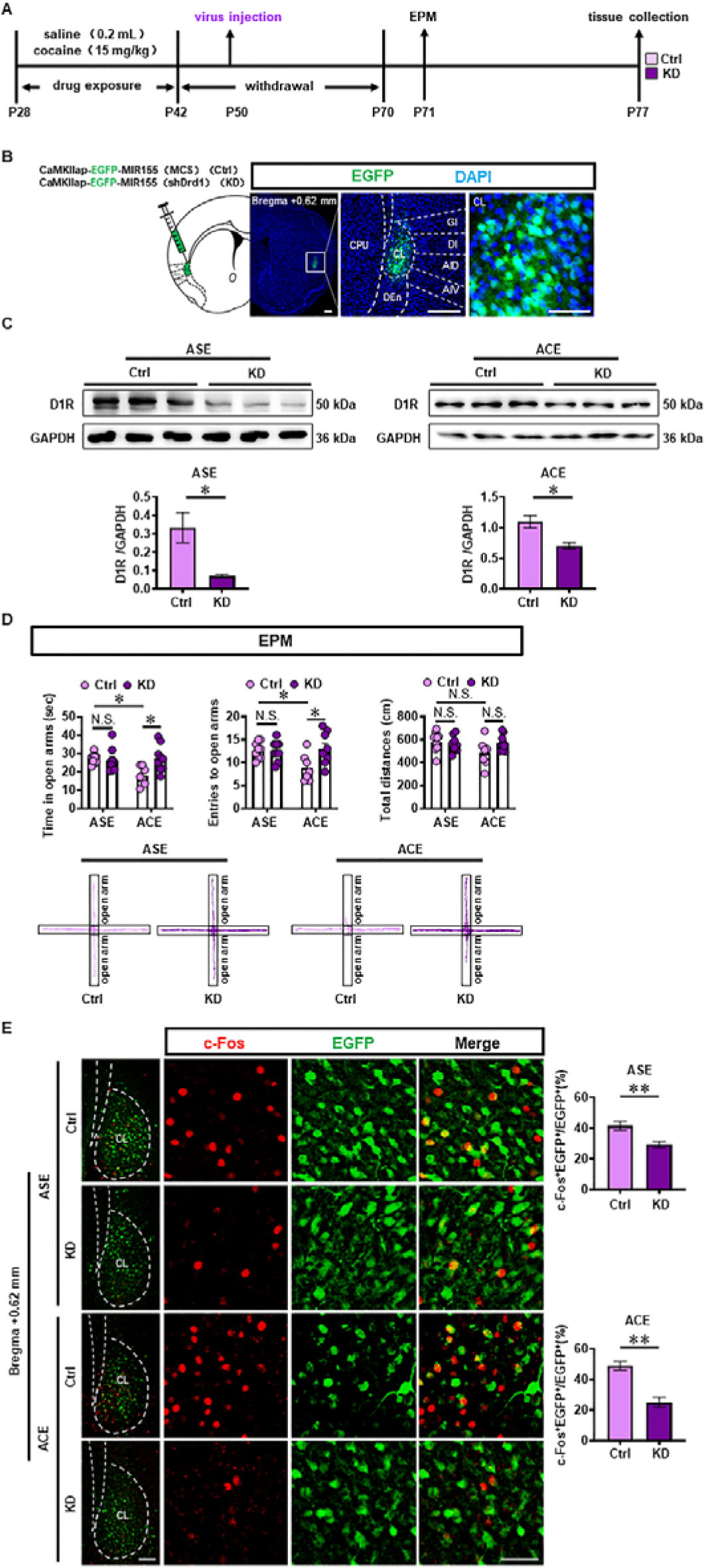
Knocking-down claustral D1R^CaMKII^ level suppresses ACE-induced anxiety-like behaviors in mice during adulthood and claustrum activation. **A**, Experimental design and timeline for KD virus treatment. **B**, Local injections of virus into claustrum. Scale bar, 300 μm. **C**, The D1R protein level. n = 3 mice per group. **D**, EPM test. Control virus-treated ASE mice, n = 9 mice; Knocking-down virus-treated ASE mice, n = 9 mice; Control virus-treated ACE mice, n = 7 mice; Knocking-down-treated ACE mice, n = 9 mice. **E**, The percentage of c-Fos-positive neurons on EGFP-transfected neurons. Control virus-treated ASE mice, n = 6 mice; Knocking-down virus-treated ASE mice, n = 6 mice; Control virus-treated ACE mice, n = 4 mice; Knocking-down-treated ACE mice, n = 6 mice. Scale bar, 100 μm/50 μm. Ctrl, control virus-treated mice; KD, knocking-down virus-treated mice; CL, claustrum. Data are presented as the Mean ± SEM. N.S., *p* > 0.05, *, *p* < 0.05, **, *p* < 0.01 vs Ctrl.

As shown in Figure 3D, Ctrl-treated ACE mice (n = 7 mice) exhibited less time spent (*t* = 3.254, *p* = 0.0168) and less entries (*t* = 2.877, *p* = 0.0431) into the open arms, while similar total distances traveled (*t* = 2.285, *p* = 0.1647) in EPM apparatus than those in Ctrl-treated ASE mice (n = 9 mice) during adulthood. When compared to Ctrl-treated ASE mice (n = 9 mice), KD-treated ASE mice (n = 9 mice) did not affect the time spent in open arms (*t* = 0.3836, *p* = 0.9993), the entries into the open arms (*t* = 0.08971, *p* >0.9999) and the total distances traveled (*t* = 0.5307, *p* = 0.9959) in EPM apparatus during adulthood. However, as shown in Figure 3D, KD-treated ACE mice (n = 9 mice) exhibited more time spent (*t* = 3.329, *p* = 0.0138) and more entries into the open arms (*t* = 3.129, *p* = 0.0231), while did not affect the total distances traveled (*t* = 2.000, *p* = 0.2863) in EPM apparatus during adulthood, when compared with Ctrl-treated ACE mice (n = 7 mice). These results indicate that knocking-down claustrum D1R^CaMKII^ could suppress ACE-induced anxiety-like behaviors during adulthood.

As shown in Figure 3E, the percentage of c-Fos-positive neurons on EGFP-transfected neurons in KD-treated ASE mice (n = 6 mice, *t* = 3.490, *p* = 0.0058) and KD-treated ACE mice (n = 6 mice, *t* = 5.297, *p* = 0.0007) were much lower than those in Ctrl-treated ASE mice (n = 6 mice) and Ctrl-treated ACE mice (n = 4 mice), respectively. These results indicate that knocking-down claustrum D1R^CaMKII^ level could decrease the activity of CaMKII-positive neurons in claustrum.

### EA treatment rescues anxiety-like behaviors, the activated claustrum activation and the increased D1R^CaMKII^ in ACE mice during adulthood

EA treatments at acupoints of GV20 and GV29 during P42-P70 were carried out in ACE mice (EA, n = 10 mice), and placebo acupoints were used as sham EA treatment (SEA, n = 13 mice) (Figure 4A, 4B). Compared with SEA-treated ACE mice, EA-treated ACE mice spent more time (*t* = 2.084, *p* = 0.0496), similar entries into the open arms (*t* = 1.536, *p* = 0.1394) and similar total distances traveled (*t* = 0.7660, *p* = 0.4522, Figure 4C) in EPM apparatus during adulthood. These results indicate that EA at GV20 and GV29 efficiently alleviate ACE-induced anxiety-like behaviors during adulthood.

**Figure 4.**
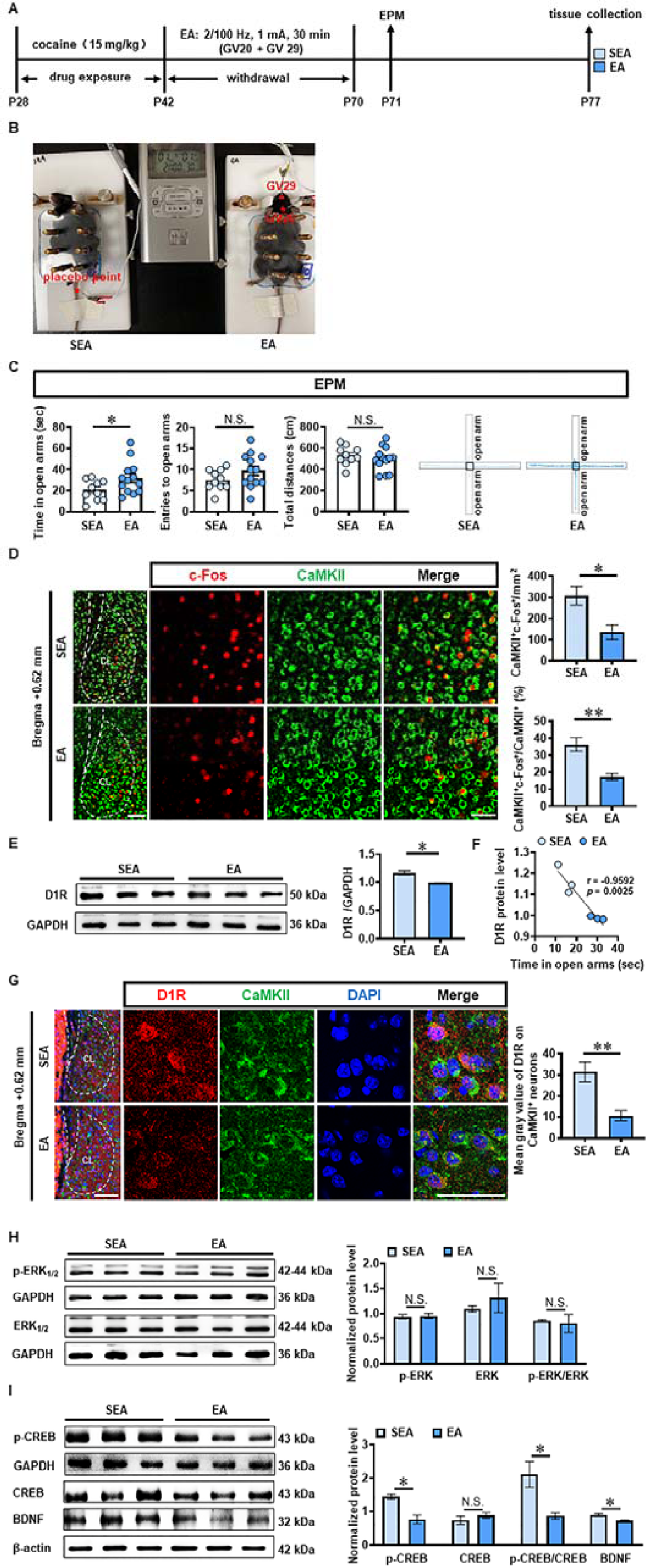
EA treatment rescues ACE-increased anxiety-like behaviors, claustrum activation and D1R^CaMKII^ level during adulthood. **A**, Experimental design and timeline for EA treatment. **B**, Schematic diagram of EA and SEA treatments in ACE mice. **C**, EPM test. EA group, n = 10 mice; SEA group, n = 13 mice. **D**, The number and percentage of c-Fos-positive & CaMKII-positive neurons. n = 6 mice per group. Scale bar, 100 μm/50 μm. **E**, The D1R protein level. n = 3 mice per group. **F**, The expression of D1R^CaMKII^. N = 6 mice per group. Scale bar, 100 μm/50 μm. **G**, The correlation analysis of claustrum D1R level with EPM data. n = 6 mice. **H**, Levels of p-ERK_1/2_ and ERK_1/2_. **I**, Levels of p-CREB, CREB and BDNF. n = 3 mice per group. SEA, sham electro-acupuncture-treated ACE mice; EA, electro-acupuncture (GV20 and GV29)-treated ACE mice; CL, claustrum. Data are presented as the Mean ± SEM. N.S., *p* > 0.05, *, *p* < 0.05, **, *p* < 0.01 vs SEA.

Compared with SEA-treated ACE mice (n = 6 mice, Figure 4D), both the number (*t* = 3.076, *p* = 0.0117) and the percentage (*t* = 4.246, *p* = 0.0017) of c-Fos-positive & CaMKII-positive neurons were much lower in EA-treated ACE mice (n = 6 mice). In parallel, the total level of claustrum D1R protein (n = 3 mice per group, *t* = 4.396, *p* = 0.0117, Figure 4E) was lower in EA-treated ACE mice than that in SEA-treated ACE mice. The claustrum D1R level was negatively correlated with the spending time in open arms of EPM apparatus in SEA-treated and EA-treated ACE mice during adulthood (n = 6 mice, *r* = -0.9592, *p* = 0.0025, Figure 4F). As shown in Figure 4G, the mean gray value of D1R on CaMKII-positive neurons was much lower (n = 6 mice per group, *t* = 3.989, *p* = 0.0026) in EA-treated ACE mice than that in SEA-treated ACE miceduring adulthood. These results indicate that EA at GV20 and GV29 could suppress CaMKII-positive neuron activity and D1R^CaMKII^ level in claustrum.

As shown in Figure 4H-J, there were no differences on protein levels p-ERK_1/2_ (*t* = 0.2242, *p* = 0.8336), ERK_1/2_ (*t* = 0.7388, *p* = 0.5010), CREB (*t* = 0.9656 *p* = 0.3889) and ratio of p-ERK_1/2_ to ERK_1/2_ (*t* = 0.3008, *p* = 0.7786) in claustrum between EA-treated ACE mice (n = 3 mice) and SEA-treated ACE mice (n = 3 mice). While, the levels of p-CREB (*t* = 4.464, *p* = 0.0111), BDNF (*t* = 2.957, *p* = 0.0417) and radio of p-CREB to CREB (*t* = 3.191, *p* = 0.0332) were lower in EA-treated ACE mice (n = 3 mice), when compared with SEA-treated ACE mice (n = 3 mice). These results imply that p-CREB and BDNF might be downstream signals of D1R targeted by EA treatment.

## Discussion

The current study explored the role of claustrum in ACE-induced anxiety during adulthood. We found that ACE produces obvious anxiety in male mice during adulthood, along with being evoked CaMKII-positive neurons and increased D1R^CaMKII^ in claustrum. Locally inhibiting D1R function or knocking-down D1R^CaMKII^ levels within claustrum efficiently suppresses ACE-increased claustrum activation, and ultimately rescues ACE-induced anxiety-like behaviors in mice during adulthood. EA at acupoints of GV20 and GV29 alleviates ACE-induce anxiety-like behaviors, potentially by targeting claustrum D1R^CaMKII^ (Figure 5).

**Figure 5.**
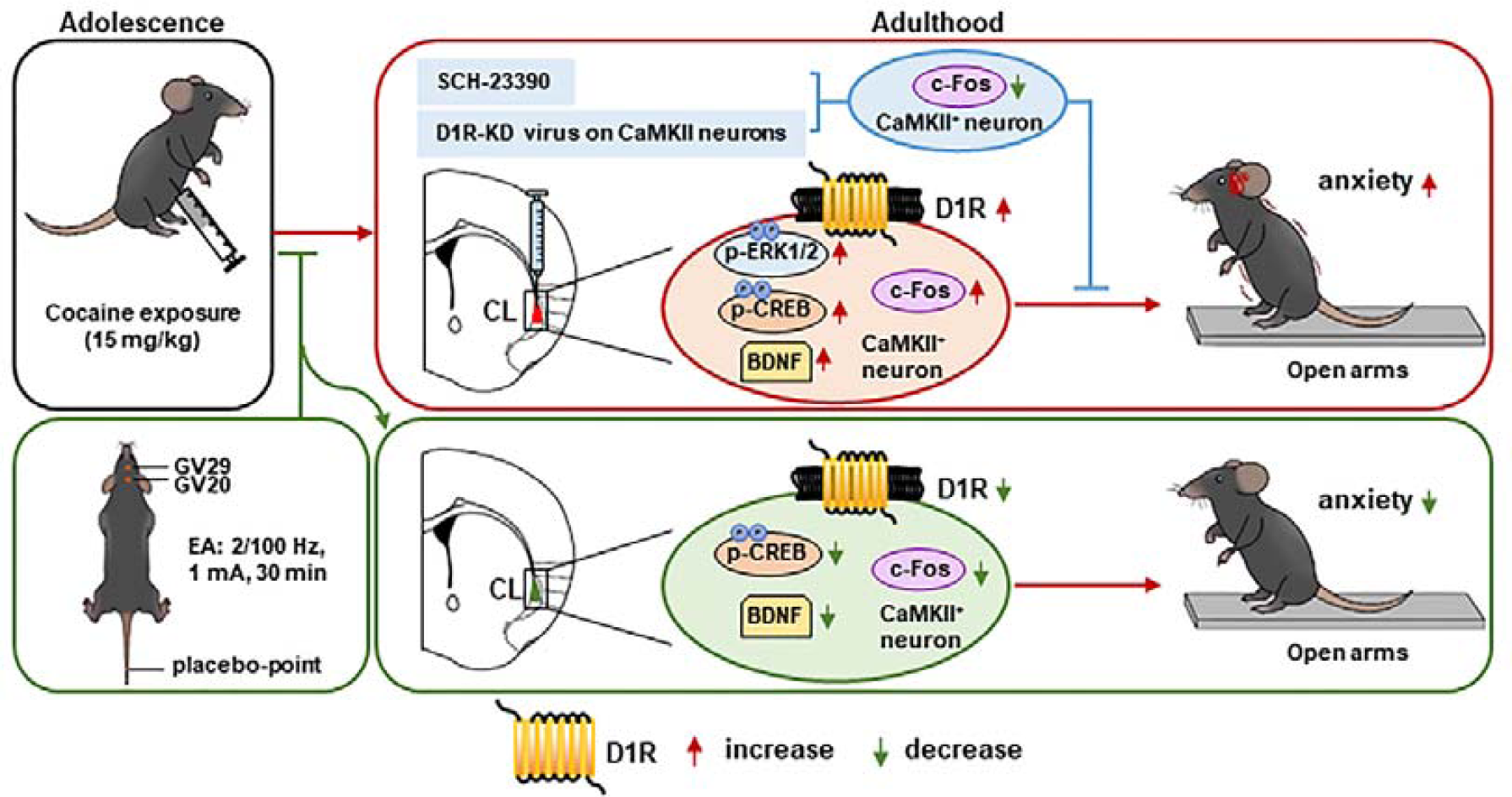
Schematic summary of present study. Adolescent cocaine exposure (ACE) enhances anxiety-like behaviors in male mice during adulthood, accompanied with triggered activation of CaMKII-positive neurons and more dopamine receptor 1 (D1R) on CaMKII-positive neurons (D1R^CaMKII^) in claustrum. Either blocking D1R with SCH-23390 or knocking-down D1R^CaMKII^ with virus in the claustrum efficiently reduces claustrum activation, and ultimately suppresses ACE-induced anxiety-like behaviors in mice. By targeting claustrum D1R^CaMKII^, electro-acupuncture (EA) at acupoints of GV20 and GV29 rescues ACE-increased anxiety-like behaviors in mice during adulthood. In the process, the p-CREB and BDNF may be the potential subsequent signals to D1R^CaMKII^.

Consistent with human and animal studies on ACE, including ours, here we found that ACE mice exhibited more anxiety-like behaviors in their adulthood. Further, we found that CaMKII-positive neurons in claustrum seem to be involved in the process. Recently, Niu et al. [29] report that claustrum is crucial encoding stress-induced anxiety, as indicated by attenuation of stress-induced anxiety with silencing CaMKII-positive neurons in claustrum. Claustrum receives inputs and sends back axonal outputs to the prefrontal cortex (PFC), a critical region coding cocaine-related anxiety [38,39]. In our previous study, we found that ACE altered the function of local circuit from GABAergic interneuron to pyramidal neurons in mPFC, which mediates ACE-increased anxiety-like behaviors in adult mice [7]. Due to the robust communications between mPFC and claustrum, ACE-triggered claustrum should contribute to the anxiety-like behaviors in mice of adulthood. Firstly, we explored the potential molecules that underlie the evoked claustrum by ACE. A recent study by Terem et al. [28] found that D1R-positive (D1R^+^) neurons in claustrum, especially those connected with PFC are required for acquisition and reinstatement of cocaine preference. They focus the cell subgroups of D1R^+^ and D1R^-^ in cocaine-preferred behaviors. Here, our findings show that it is D1R^CaMKII^ that were elevated by ACE in claustrum of adult mice, and we focus on the role of D1R in activation of CaMKII-positive neurons and ACE-related anxiety-like behaviors. Moreover, locally blocking D1R or knocking-down D1R^CaMKII^ in claustrum reduced ACE-evoked claustrum and subsequent anxiety-like behaviors. These findings indicate that activity of claustrum CaMKII-positive neurons encodes ACE-increased anxiety-like behaviors, which is mediated at least partially by D1R^CaMKII^. In other D1R-rich brain regions, such as PFC and hippocampus, cocaine abuse could alter p-ERK [40], p-CREB [41] and BDNF [42,43]. Here, we found that protein levels of p-ERK, p-CREB and BDNF were significantly increased in claustrum of ACE mice, implying that they are potential downstream signals for claustral D1R^CaMKII^ in the process of ACE-induced anxiety-like behaviors.

At present, neither D1R antagonists nor specific virus-regulated D1R could be clinically used to treat emotional disorders such as ACE-induced anxiety in humans. EA has been widely applied in management of substance abuse, especially for drugs-caused withdrawal symptoms, including anxiety [31,33,34]. Shao et al [44] reported that EA at Zusanli (ST36) and Kunlun (BL60) could reduce anxiety-like behaviors in animals by suppressing neuropeptide Y in anterior cingulate cortex. Wu et al. [45] found that EA at ST36 and Sanyinjiao (SP6) alleviated anxiety by activating D1R or antagonizing D2R in the basolateral amygdala. Previously, we found that EA with a stimulus of a mixed frequency of 2/100 Hz could alleviate ACE-induced anxiety-like behaviors in mice [34]. With the findings of claustrum here, we speculated that claustrum D1R^CaMKII^ might be one pharmacological targets of EA treatment. As expected, we found that EA at acupoints of GV20 and GV29 reversed ACE-enhanced claustrum activation by suppressing claustral D1R^CaMKII^, which ultimately ameliorated ACE-induced anxiety-like behaviors during adulthood.

## Conclusions

Our findings identified a novel role of claustrum in ACE-induced anxiety-like behaviors, and put new insight into the function of claustrum D1R^CaMKII^. EA treatments rescue ACE-induced anxiety-like behaviors potentially by targeting claustrum D1R^CaMKII^. The claustrum D1R^CaMKII^ might be a promising pharmacological target to treat drugs-induced anxiety-like behaviors.

## Abbreviations

ACE: adolescent cocaine exposure; ASE: adolescent saline exposure; Ctrl, control; D1R: dopamine receptor 1; D1R^CaMKII^, dopamine receptor 1 on CaMKII-positive neuron; D2R: dopamine receptor 2; D3R: dopamine receptor 3; EA: Electro-acupuncture; EPM: Elevated plus maze; KD, knocking-down; SEA: Sham electro-acupuncture; N.S., no significance.

## Author contributions

Chen L, Liu Z and Zhao Z have equal contribution to the manuscript. Chen L, Fan Y, Liu Z and Zhao Z: Methodology, Investigation. Du D, Pan W, Wei X and Nie J: Data Curation. Fan Y, Ge F and Ding J: Data Curation. Guan X and Fan Y: Formal analysis, Writing - Original Draft, Writing - Review & Editing. Kim HY: Writing - Review & Editing. Guan X: Supervision, Conceptualization, Project administration.

## Data availability statement

The data that support the findings of this study are available from the corresponding author upon reasonable request.

